# Culture media, DMSO and efflux affect the antibacterial activity of cisplatin and oxaliplatin

**DOI:** 10.1101/2022.03.14.484244

**Authors:** Arya Gupta, Lorenzo Bernacchia, Neil M. Kad

## Abstract

Cisplatin was originally discovered through its antibacterial action, and subsequently has found use as a potent broad spectrum anticancer agent. This study determines the effect of growth media and solvent on the antibacterial activity of cisplatin and its analogue, oxaliplatin. *E. coli* MG1655 or MG1655 *ΔtolC* were treated with the platinum compounds under different conditions and susceptibility was determined. Our results showed that DMSO reduced the activity of cisplatin by 4-fold (MIC 12.5 mg/L) compared with 0.9% NaCl-solubilized cisplatin (MIC 3.12 mg/L) when tested in MOPS. Surprisingly, complete loss of activity was observed in Mueller Hinton Broth II (MHB II). By supplementing MOPS with individual components of MHB II such as the sulphur containing amino acids, L-cysteine and L-methionine, individually or in combination reduced activity by ≥8-fold (MIC ≥25 mg/L). Oxaliplatin was less active against *E. coli* (MIC 100 mg/L) but exhibited similar inactivation in the presence of DMSO, MHBII or MOPS spiked with L-cysteine and L-methionine (MIC ≥400 mg/L). Our data suggest that the antibacterial activity of cisplatin and oxaliplatin is modulated by both choice of solvent and composition of growth media. We demonstrate that this is primarily due to sulphur-containing amino acids cysteine and methionine, an essential component of the recommended media for testing antimicrobial susceptibility, MHBII.

**Significance and impact of the study:** As well as an anticancer treatment, cisplatin possesses antibacterial activity and is active against AMR resistant persister cells, opening the possibility of renewed use against resistant bacterial strains. Our findings provide evidence on how the composition of growth media and choice of solvent modulate the antibacterial activity of cisplatin and its analogue, oxaliplatin. These observations provide a necessary, consistent standard for assessing the antibacterial activity of platinum-based compounds, as a precursor towards their application against bacterial infection.

**Graphical abstract:** 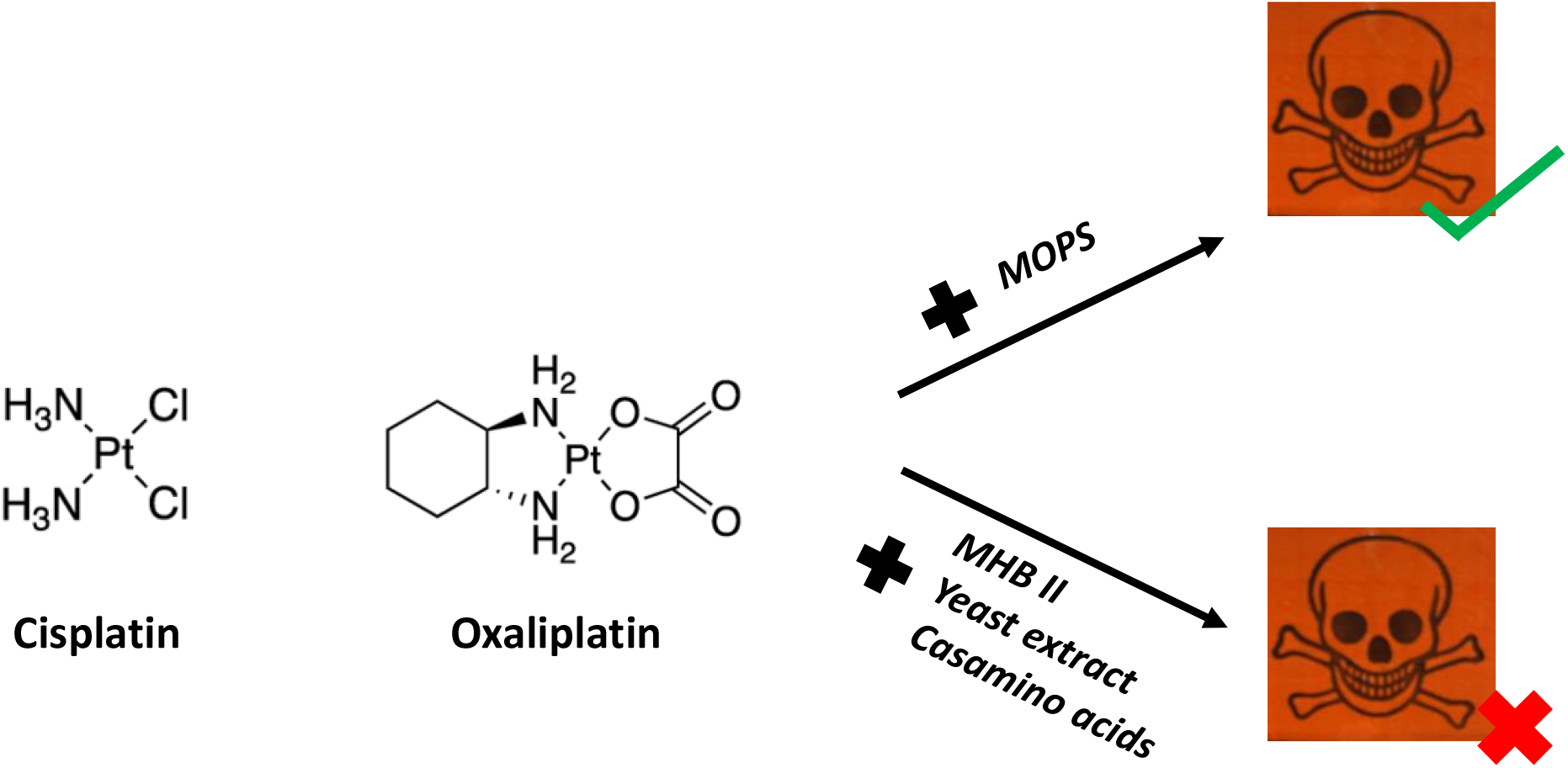

## Introduction

Cisplatin is a platinum-based DNA crosslinker mainly used to treat a variety of cancers such as testicular, ovarian, bladder, and lung (Dasari and Tchounwou 2014). It exhibits antitumor activity by forming intra- and inter-strand DNA crosslinks (Hashimoto *et al*. 2016). Although widely used in cancer chemotherapy, it was first discovered for its antibacterial activity against Gram-negative bacteria (Rosenberg *et al*. 1967); and has been shown to be effective against bacterial persister cells (Choudhury *et al*. 2016). The antibacterial activity was shown to be caused by the formation of DNA adducts through susceptibility studies using nucleotide excision DNA repair mutant strains (Beck *et al*. 1985). More recently it has been shown that genes involved in nucleotide excision repair and SOS response in *E. coli* are upregulated upon cisplatin treatment, providing further evidence of DNA damage in bacteria (Beaufay *et al*. 2020). Although a highly potent chemotherapeutic agent, cisplatin exhibits low solubility, and its activity is dependent on solvent interactions. The universal solvent, DMSO has previously been shown to reduce the cytotoxicity of cisplatin against thryocytes and stage IIb appendicular osteosarcoma in dogs (Massart *et al*. 1993; Dernell *et al*. 1998). Here we sought to establish whether both media composition and DMSO affects the antimicrobial activity of cisplatin against *E. coli*. This study also aimed to establish whether it is a substrate for the efflux pump, TolC. Determining the correct solution conditions to study cisplatin or oxaliplatin is an essential first step for reproducible testing of bacteria using these compounds.

## Results and Discussion

Cisplatin binds to DNA in both pro- and eukaryotic cells inhibiting essential processes such as transcription and replication. This DNA binding requires cisplatin to be ‘aquated’ in the cells by the replacement of a chloride ligand with water (Hall *et al*. 2014). Our initial studies of cisplatin’s antibacterial activity against *E. coli* MG1655 in MHBII showed no growth inhibition (MIC >50 mg/L, Figure 1A). In contrast, in MOPS, cisplatin exhibited a MIC of 3.12 mg/L and 0.39 mg/L, against MG1655 and MG1655 Δ*uvrA*, respectively (Figure 1A). We also sought to determine whether DMSO affected the inhibitory activity of cisplatin against *E. coli*. The MIC for DMSO was determined to be 10% (Figure 1B), therefore 2.5% was selected as the working concentration to avoid any impact on bacterial growth. Even with a low concentration of DMSO (2.5%), we found that the activity of cisplatin was reduced 4-fold and 8-fold against MG1655 and Δ*uvrA*, respectively (Figure 1A; MIC, 12.5 mg/L and 3.12 mg/L). A similar 4-fold reduction in cisplatin activity was also observed when cisplatin was dissolved in 100% DMSO instead of 0.9 % NaCl (Figure 1C). Previous studies have shown that sulphur containing compounds, such as DMSO, can bind prior to aquation and impair cytotoxicity (Hall *et al*. 2014; Fischer *et al*. 2008; Yi and Bae 2011). Consistent with these previous observations, we show for the first time that DMSO also reduces cisplatin toxicity in *E. coli* and define the MIC with 2.5% DMSO clearly as 12.5 mg/L (Figure 1A).

**Figure 1:**
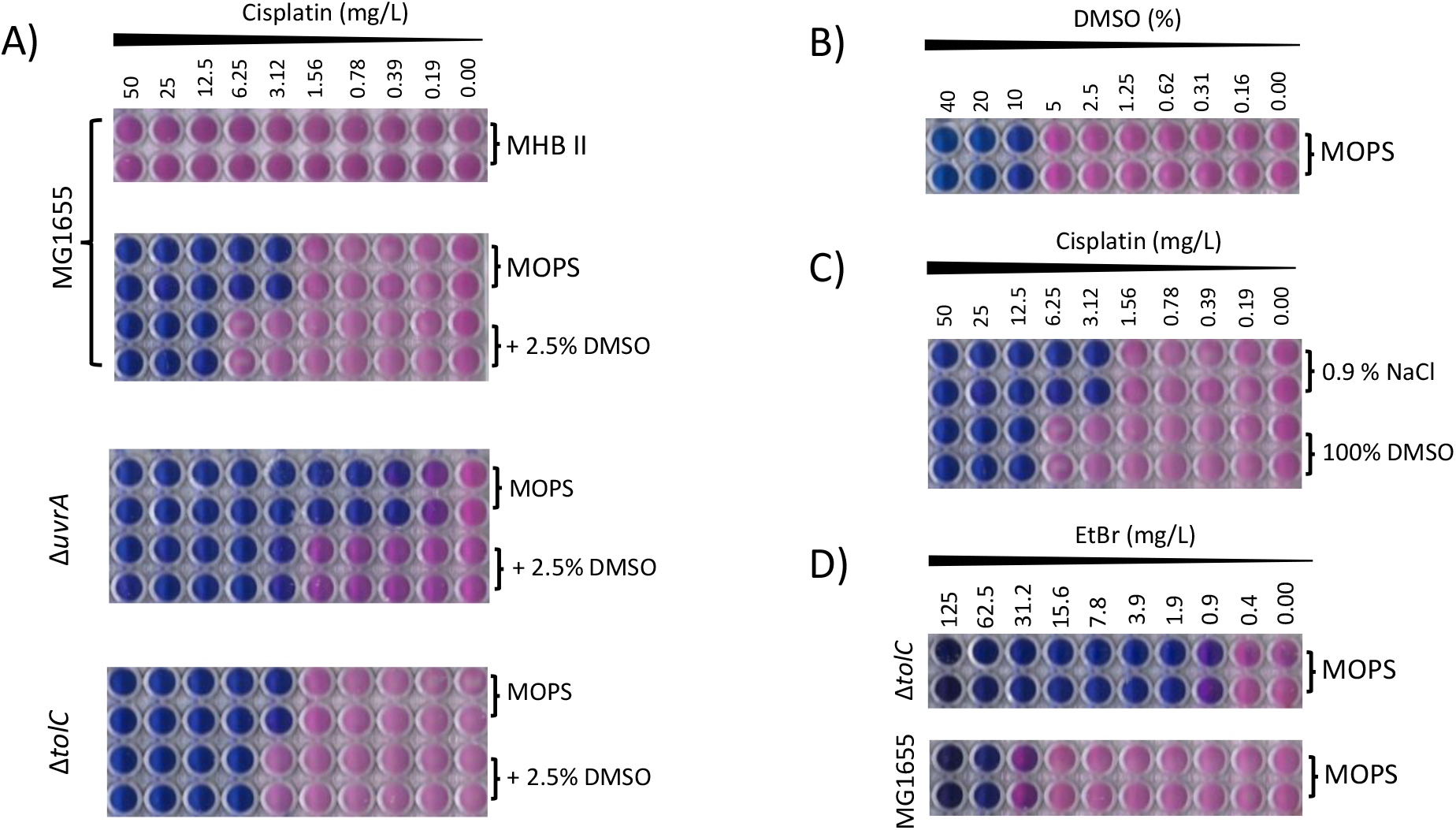
Reduced antibacterial activity of cisplatin in MHBII and in the presence of DMSO. **A)** Antimicrobial activity of cisplatin (CPt) dissolved in 0.9% NaCl in MHBII and MOPS (with and without DMSO) against MG1655, MG1655 ΔtolC and MG1655 ΔuvrA; **B)** DMSO MIC against MG1655, **C)** Inhibitory activity of cisplatin when dissolved in 0.9% NaCl or 100% DMSO, **D)** Characterisation of MG1655 ΔtolC by determining the MIC for Ethidium bromide (EtBr); (n≥3, individual plate photographs are representative of 3 independent replicates)

To eliminate efflux of cisplatin as the cause of this reduced activity, the *tolC* deletion from the Keio collection was transduced into MG1655 to generate the strain MG1655 Δ*tolC*. The susceptibility of this strain to cisplatin was identical to WT (MIC 3.12 mg/L, Figure 1A), compared with a 64-fold difference observed for ethidium bromide inhibitory activity, against WT and Δ*tolC* (MIC, 62.5 mg/L and 0.9 mg/L, respectively, Figure 1D). Ethidium bromide was used as a control, since it is a known substrate for TolC that possesses DNA intercalation activity (Paixao *et al*. 2009). Our findings clearly demonstrate that cisplatin is not a substrate for the AcrAB-TolC efflux pump since its MIC was not altered when we knocked out TolC, an integral part to the system.

The primary differences between MHBII and minimal MOPS are the presence of casamino acids and beef extract, at concentrations of 17.5 g/L and 3 g/L, respectively. To test if these components were responsible for the elimination of antibacterial activity in MHBII, we determined the activity of cisplatin in MOPS with the addition of either casamino acids (17.5 g/L) or yeast extract (3 g/L) (used in place of beef extract). Figure 2A shows the antibacterial activity of cisplatin was markedly diminished upon the addition of the two media supplements. To confirm this observation is specific to cisplatin, we determined the MIC for ampicillin and nadifloxacin (bactericidal and bacteriostatic compounds, respectively), in MOPS and MOPS supplemented with either casamino acids or yeast extract. The largest change observed was ~4-fold, well within EUCAST breakpoints (EUCAST, version 8.1.2018), and considerably smaller than that observed with cisplatin in similar conditions (Figure 2A). Therefore, the small increase in MIC observed in the case of ampicillin is not significant and shows that growth does not play a role in the observed cisplatin susceptibility.

**Figure 2:**
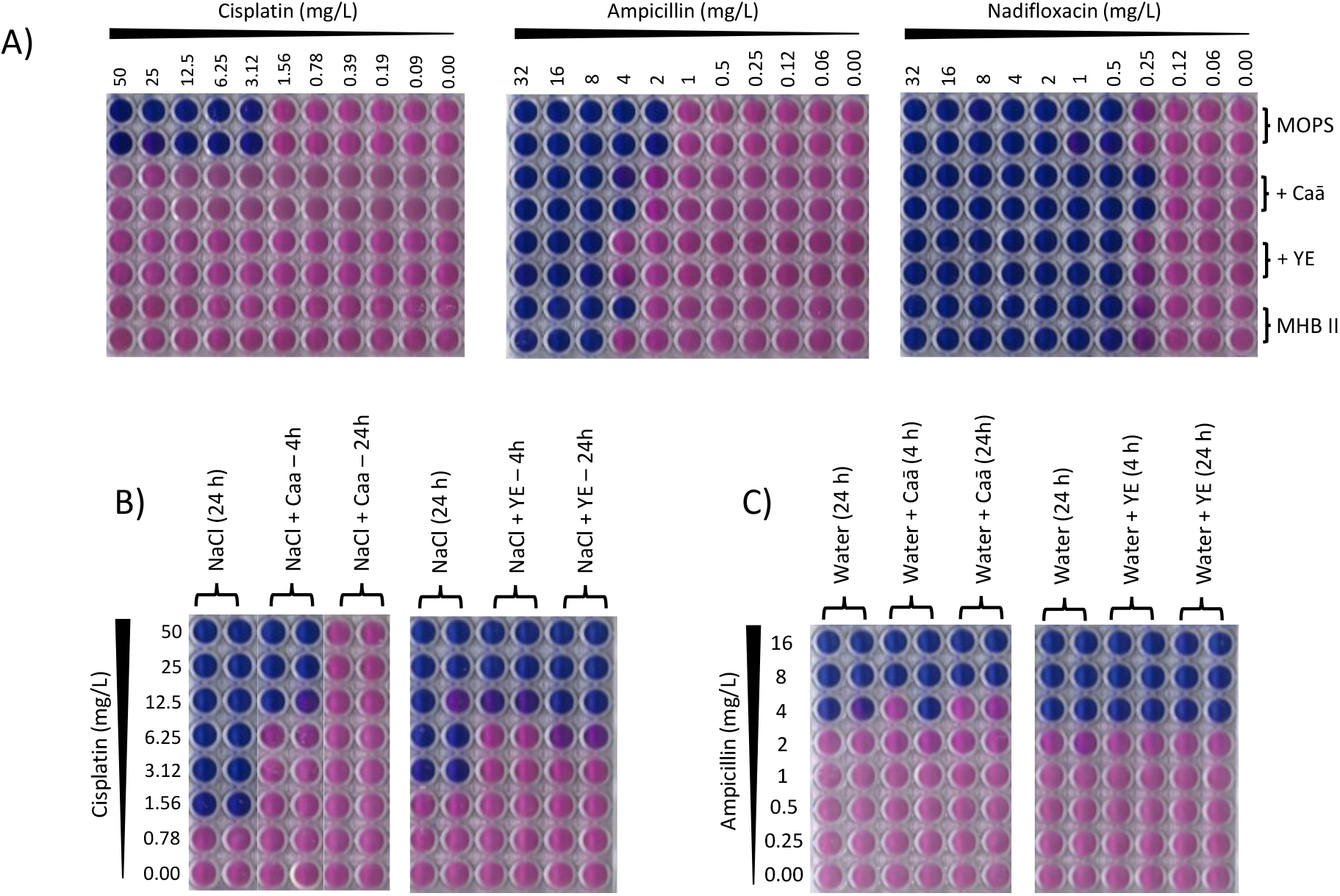
Role of growth media composition in the reduced susceptibility of cisplatin. **A)** Inhibitory profile of Cisplatin (CPt), ampicillin (Amp) and nadifloxacin against MG1655 in MOPS, MOPS supplemented with either 17.5 g/L casamino acids (Caa) or 3 g/L yeast extract (YE) and MHBII; **B)** Activity profile of CPt in MOPS after incubation with either 17.5 g/L Caa or 3 g/L YE in 0.9% NaCl for 4 h and 24 h; **C)** Inhibitory activity of Amp in MOPS after incubation with either Caā or YE in water; (n≥3, individual plate photographs are representative of 3 independent replicates)

To further test if the reduced activity is not dependent on growth but rather a direct interaction between cisplatin and the components of MHBII, we preincubated cisplatin in 0.9% NaCl with either casamino acids (acid hydrolysate of casein, 17.5 g/L) or yeast extract (3 g/L), for 4 h and 24 h, and measured the MIC. As a control, ampicillin was subjected to the same treatment in water. An 8-fold and >16-fold reduction in activity was observed when cisplatin was incubated with casamino acids (Figure 2B). Cisplatin activity was also reduced by 8-fold, upon treatment with yeast extract (Figure 2B). Neither of the supplements influenced the activity of ampicillin (Figure 2C), suggesting that the reduced activity of cisplatin is not linked to growth deficiency. Rather, cisplatin likely interacts with methionine and/or cysteine, found in the acid hydrolysate of casein (casamino acids) used in MHBII, thereby reducing its availability and subsequent activity. Because acid treatment of cysteine results in numerous products, we simplified our approach by studying cisplatin in the presence of minimal MOPS supplemented with L-cysteine (4.6 g/kg) and L-methionine (32 g/kg), which reflect the amounts estimated to be present upon acid hydrolysis of casein (Pieniazek *et al*. 1974). In these conditions, we observed a significant reduction in cisplatin activity (MIC, ≥25 mg/L, Figure 3A). As in Figure 2A, ampicillin exhibited a 4-fold reduction in activity under the same conditions (Figure 3B). We expect that the sulphur present in cysteine and methionine react with one of cisplatin’s platinum atoms, leading to its inability to bind to DNA, akin to DMSO (Hall *et al*. 2014). Our results also correspond to the antagonistic role of methionine and cysteine on the antibacterial activity of cisplatin shown previously in *Helicobacter pylori* (Lettl *et al*. 2020).

**Figure 3:**
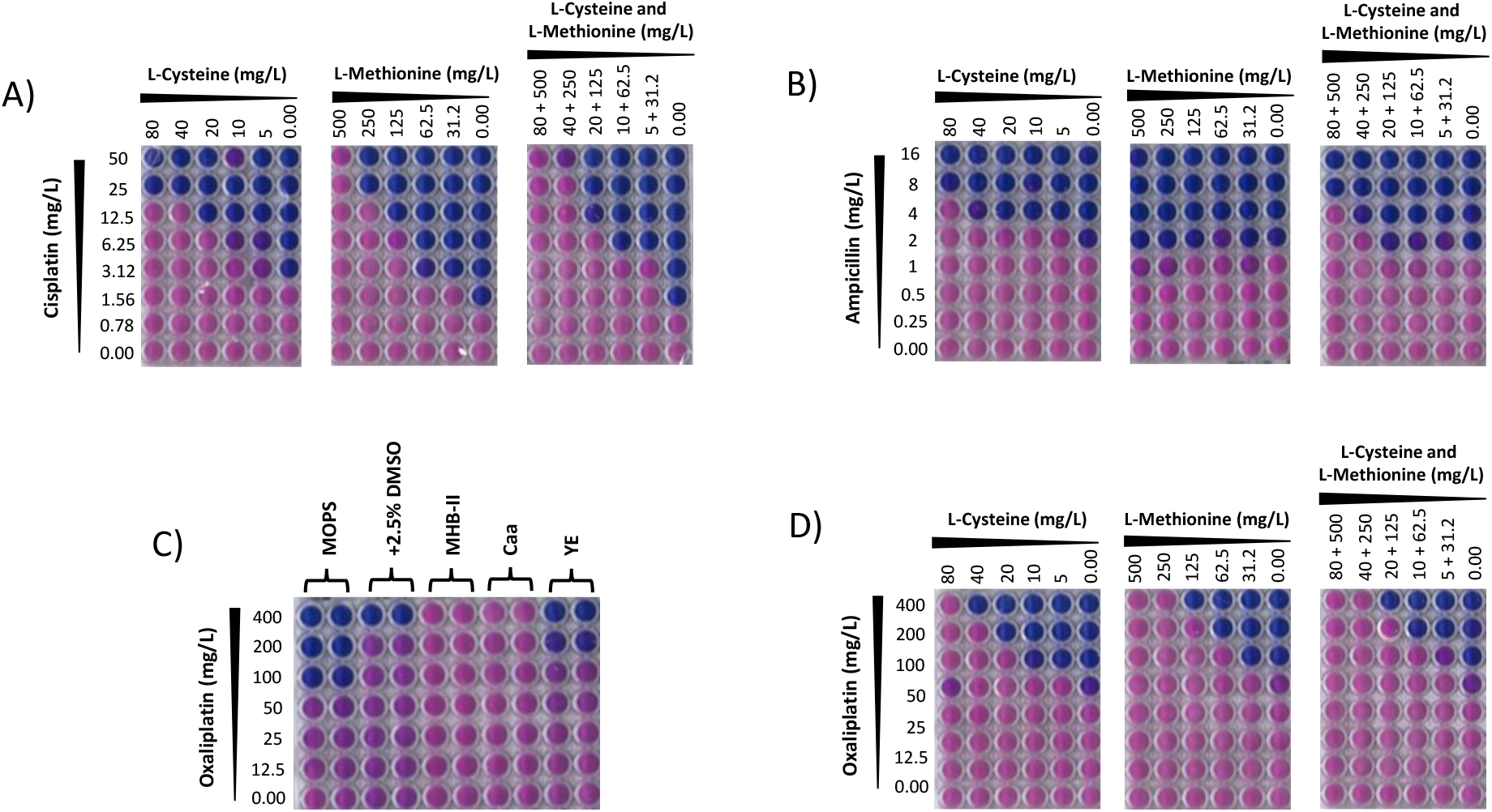
Reduced antibacterial activity of platinum drugs, cisplatin and oxaliplatin in the presence of sulphur containing amino acids. **A)** Cisplatin checkerboard assays with L-cysteine and L-methionine, individually and together; **B)** Ampicillin checkerboard assays with L-cysteine and L-methionine, individually and together; **C)** Antibacterial activity of oxaliplatin against MG1655 in MOPS, MHBII and MOPS treated with components of MHBII; **D)** Oxaliplatin checkerboards with L-cysteine and L-methionine. (n≥3, individual plate photographs are representative of at least 3 independent replicates) We also show that the reduction in antibacterial activity in MHBII is not specific to cisplatin. Figure 3C shows that oxaliplatin (a structural analogue of cisplatin), was less active against MG1655 (MIC 100 mg/L, Figure 3C) compared with cisplatin. Furthermore, oxaliplatin shows no activity in MHBII (MIC >400 mg/L), and reduced activity in the presence of L-cysteine and L-methionine individually or in combination (MIC >400 mg/L, Figure 3D).

In summary, our results explain the necessity for high cisplatin concentrations used in previous studies (Choudhury *et al*. 2016; Beck *et al*. 1985; Keller *et al*, 2001), and provide a framework for the accurate study of cisplatin-based bacterial cytotoxicity for future studies. We recommend the use of minimal media with limited DMSO concentrations of <2.5%. Using the conditions laid out in this study, we hope to provide a uniform standard for studying bacterial cytotoxicity with platinum-based compounds such as cisplatin and oxaliplatin.

## Material and methods

### Bacterial strains, media culture conditions

The strains used in this study include *E. coli* MG1655, MG1655 Δ*uvrA*, and MG1655 Δ*tolC*. The two knockout strains were generated by P1 transduction of the respective gene deletions from the Keio collection (Baba *et al*. 2006). Luria Bertani broth and agar were used to maintain bacterial strains (Sigma, Dorset, UK). Strains were sub-cultured in Mueller Hinton Broth II (MHBII, Sigma, Dorset, UK) or MOPS (Melford, Berkshire, UK) minimal medium (Neidhart *et al*. 1974) supplemented with 0.2% glucose, 1.32 mM K_2_HPO_4_ and 0.1 μg/mL thiamine, for susceptibility testing. All strains were grown at 37°C with vigorous aeration.

### Antimicrobials and chemicals

Cisplatin, oxaliplatin and nadifloxacin were purchased from Sigma, Dorset, UK, and ampicillin was sourced from Melford, Suffolk, UK. Media supplements – casamino acids and yeast extract were purchased from BD Scientific, Berkshire, UK.

### Antimicrobial susceptibility testing

The minimum inhibitory concentration (MIC) of cisplatin, oxaliplatin, ampicillin and nadifloxacin, was determined by broth microdilution against MG1655 and MG1655 Δ*uvrA*, according to CLSI guidelines (CLSI 2012), with the following adjustments. After 16 h incubation of 96-well plates at 37°C the MIC was determined by addition of 50 μL of resazurin (0.3 mg/mL) in MOPS/Tricine buffer (pH 7.8) to the plates and incubation at 37°C for a further 4 h. A colour change of blue to pink indicated growth and the MIC was defined as the lowest concentration which prevented a colour change (Palomino *et al*. 2009). The use of resazurin provides a highly accurate determination of the MIC, since even one ‘live’ cell will cause a colour change. Top stocks of cisplatin were made at 0.5 mg/L in either 0.9% NaCl or DMSO due to its low solubility and were subsequently used as the starting concentration for susceptibility testing, yielding an unconventional doubling dilution series. Obtained MICs were not rounded off to the nearest value indexed to the base 2.

## Acknowledgements

The authors would like to thank the other members of Kad group for helpful discussions on the topic.

## Funding

This work was supported by Cancer Research UK. Grant No. - [C70469/A30456]

## Transparency declarations

None to declare

## Notes

### Competing Interest Statement

The authors have declared no competing interest.

